# Interphotoreceptor matrix proteoglycans IMPG1 and IMPG2 proteolyze in the SEA domain and reveal localization mutual dependency

**DOI:** 10.1101/2022.04.27.489588

**Authors:** Benjamin Mitchell, Chloe Coulter, Werner J. Geldenhuys, Scott Rhodes, Ezequiel M. Salido

**Affiliations:** Department of Ophthalmology and Visual Sciences, West Virginia University, Morgantown, WV, USA; Undergraduate Program in Biochemistry, West Virginia University, Morgantown, WV, USA; Department of Neuroscience, School of Medicine, West Virginia University, Morgantown, WV, USA; Department of Pharmaceutical Sciences, School of Pharmacy, West Virginia University, Morgantown, WV, USA; Department of Biochemistry, West Virginia University, Morgantown, WV, USA

**Keywords:** Interphotoreceptor matrix, IMPG1, IMPG2, SEA domain, proteolysis, proteoglycan, chondroitin sulfate, retina, retinitis pigmentosa

## Abstract

The interphotoreceptor matrix (IPM) is a specialized extracellular mesh of molecules surrounding the inner and outer segments of photoreceptor neurons. Interphotoreceptor matrix proteoglycan 1 and 2 (IMPG1 and IMPG2) are major components of the IPM. Both proteoglycans possess SEA (sperm protein, enterokinase and agrin) domains, which may support proteolysis. Interestingly mutations in the SEA domains of IMPG1 and IMPG2 are associated with vision disease in humans. However, if SEA domains in IMPG molecules undergo proteolysis, and how this contributes to vision pathology is unknown. Therefore, we investigated SEA-mediated proteolysis of IMPG1 and IMPG2 and its significance to IPM physiology. Immunoblot analysis confirmed proteolysis of IMPG1 and IMPG2 in the retinas of wildtype mice. Point mutations mimicking human mutations in the SEA domain of IMPG1 that are associated with vision disease inhibited proteolysis. These findings demonstrate that proteolysis is part of the maturation of IMPG1 and IMPG2, in which deficits are associated with vision diseases. Further, immunohistochemical assays showed that proteolysis of IMPG2 generated two subunits, a membrane-attached peptide and an extracellular peptide. Notably, the extracellular portion of IMPG2 trafficked from the IPM around the inner segment toward the outer segment IPM by an IMPG1-dependent mechanism. This result provides the first evidence of a trafficking system that shuttles IMPG1 and IMPG2 from the inner to outer IPM in a co-dependent manner. In addition, these results suggest an interaction between IMPG1–IMPG2, and propose that mutations affecting one IMPG could affect the localization of the normal IMPG partner contributing to the disease mechanism of vision diseases associated with defective IMPG molecules.

## Introduction

The extracellular space between neurons and glial cells is supported by a network of macromolecules known as the extracellular matrix. In the retina, the interphotoreceptor matrix (IPM) is a specialized extracellular matrix surrounding the inner segment (IS) and outer segment (OS) of photoreceptors, extending between Müller cells and the retina pigment epithelium (RPE). The IPM is thought to have a fundamental role in transporting nutrients and metabolites between photoreceptors and RPE. Other proposed roles of the IPM are to buffer extracellular calcium and facilitate growth factor presentation and cell–cell communication ^1^.

The IPM is primarily composed of two retinal-specific proteoglycans: interphotoreceptor matrix proteoglycan 1 and 2 (IMPG1 and IMPG2), also known as SPARC and SPARCAN or IPM150 and IPM200, respectively ^2-6^. Both proteoglycans are densely glycosylated and rich in chondroitin sulfate (ChS). Our previous work shows that IMPG1 is a secreted proteoglycan homogeneously distributed throughout the retina IS and OS, while transmembrane proteoglycan IMPG2 seems to be strictly located in the IS ^7^. Interestingly, we also showed that localization of IMPG1 depends on the presence of IMPG2.

IMPG1 and IMPG2 share common protein domains termed SEA (sperm protein, enterokinase and agrin) domains ^8^. SEA domains are associated with highly glycosylated proteoglycans including the mucins family, which expresses well-studied SEA domains ^9^. IMPG1 and IMPG2 each have two SEA domains, SEA-1 and SEA-2 (Fig. 1A) ^3,4,6,10^. Missense mutations affecting the SEA-2 domain of IMPG1 are associated with development of retinitis pigmentosa (RP) in humans ^11^. RP is characterized by progressive cell death of rod photoreceptors, leading to tunnel vision and night blindness ^12^.

**Figure 1.**
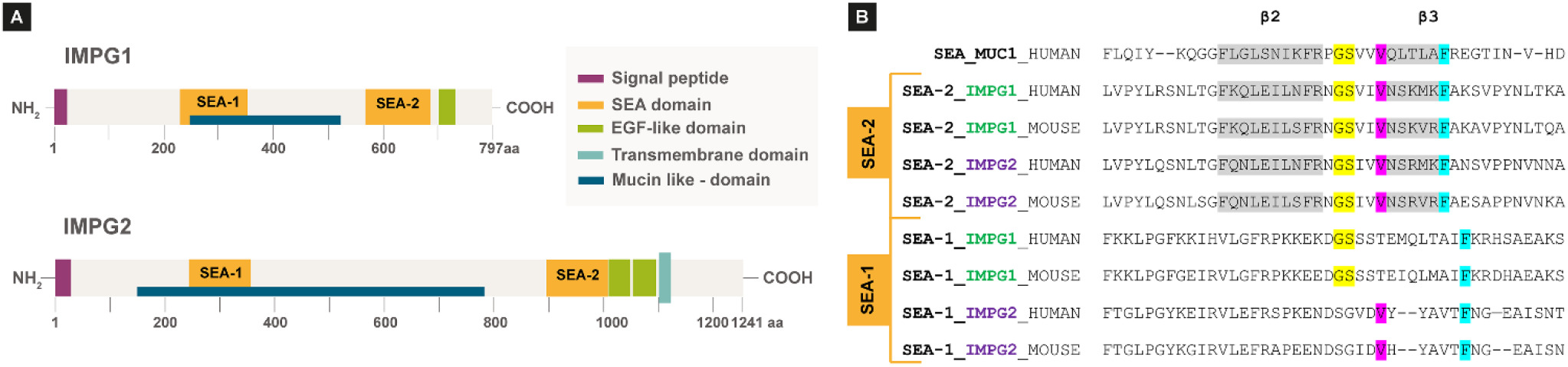
IMPG1 and IMPG2 SEA-2 domains have amino acid sequences compatible with proteolysis. **A**, Schematic of IMPG1 and IMPG2 domains. Only IMPG2 has a transmembrane domain; SEA-1 and SEA-2 are present in both proteoglycans. **B**, Amino acid alignment of known self-proteolytic SEA domain of MUC1 protein against SEA-2 and SEA-1 sequences from IMPG1 and IMPG2, in humans and mice. Established proteolytic GSxxV sequence is in yellow and violet. Conserved amino acid sequences between SEA domains are in grey and cyan.

Depending on the amino acid sequence, some mucin SEA domains undergo intramolecular proteolysis, while others do not ^13-16^. Proteolytic SEA domains possess the conserved amino acid sequence, GSxxV, where x positions are occupied by V or I; this motif is found in the SEA-2 domains of both IMPG proteins ^14,15,17^. However, whether SEA domains in IMPG molecules undergo proteolysis is unknown.

We hypothesized that IMPG1 and IMPG2 undergo proteolysis, and that mutations in IMPG1 disrupt proteolysis, causing RP. We also speculate that proteolyzed IMPG2 may have different retina localization than previously described. Because localization of IMPG1 depends on the presence of IMPG2, we also hypothesized that localization of IMPG2 may depend on IMPG1 as well. Thus, we examined the role of SEA domains in IMPG1 and IMPG2.

## Results

### Predicted SEA domain proteolysis of IMPG1 and IMPG2

Comparison of the mucin-1 SEA amino acid sequence with the SEA-1 and SEA-2 sequences of IMPG1 and IMPG2 (Fig. 1A) showed that SEA-2 domains from both IMPG proteins contain the canonical amino acid sequence for intramolecular proteolysis (Fig. 1B). Alignment also revealed that SEA-1 domains in IMPG proteins do not contain this sequence. Therefore, IMPG1 and IMPG2 have predicted capacity to proteolyze at their SEA-2 but not SEA-1 domains. Alignment of murine IMPG SEA domains revealed the same anticipated catalytic activity for the SEA-2 domain (Fig. 1B), consistent with conservation across species ^17^.

### IMPG1 undergoes intramolecular proteolysis

Both IMPG1 and IMPG2 proteoglycans are heavily glycosylated and contain ChS glycosaminoglycan chains ^4,18^. The predicted molecular weight of IMPG1 is 89 kDa, and proteolysis at the SEA-2 domain would generate protein fragments of 71 and 18 kDa (Fig. 2A). To determine the relative molecular mobility (Rf) of the IMPG1 backbone, ChS and N-/O-glycosylation were removed using a series of enzymatic digestions in isolated mouse retina tissue using chondroitinase ABC (ChSabc) and deglycosylation enzyme mix (Deglyco) (Fig. 2B). Western blot of digested proteins with an antibody recognizing the 71-kDa peptide section of IMPG1 showed a band in wildtype (WT) samples that was absent in retinas of IMPG1-knockout (KO) mice. WT samples treated with ChSabc had an Rf of ∼100 kDa; however, WT samples treated with ChSabc and Deglyco showed a ∼71-kDa band, consistent with IMPG1 proteolysis. Total protein loading controls are included in the supporting information (Fig. S1). Samples required treatment with ChSabc to be detected by the primary antibody.

**Figure 2.**
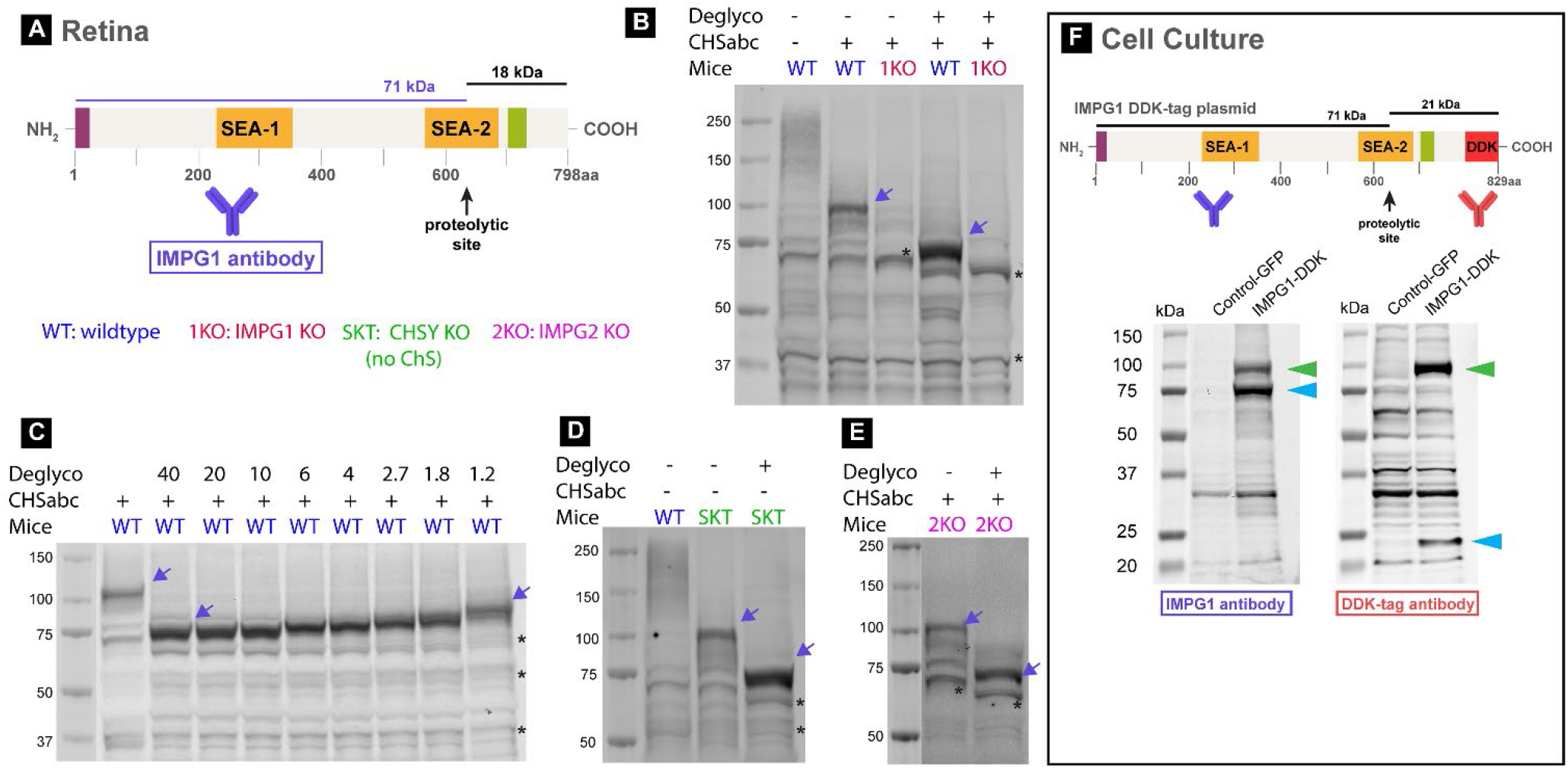
IMPG1 undergoes proteolysis. **A**, Schematic of IMPG1 structure representing the two expected peptides after proteolysis and respective relative molecular mobility (Rf). **B**, Western blot assay using retina samples from wildtype (WT) and IMPG1 KO mice (1KO), treated with and without chondroitinase abc (ChSabc) as well as with and without deglycosylation enzymes (Deglyco). Membranes were stained with an antibody against the long IMPG1 predicted peptide (71 kDa) (violet arrows). WT samples co-treated with ChSabc and Deglyco enzymes yielded one single IMPG1 band with an Rf near 71 kDa. Black asterisks mark nonspecific bands. **C**, WT samples treated with ChSabc, exposed to a series of decreasing amounts of Dglyco enzyme (nL/μL of sample). Only one single IMPG1 band is present in all serial dilutions, and its molecular weight increases stepwise with decreasing Dglyco enzyme. **D, E**, Molecular weight profile of IMPG1 in a mouse model lacking chondroitin sulfate (SKT mice) (**D**) and in IMPG2 KO mice (2KO) (**E**). **F**, Western blot assay of HEK293 cell samples expressing GFP as a negative control or IMPG1-DDK transgenes. Antibodies against IMPG1 or DDK-tag were used in the same membrane. Light blue arrows indicate long and short IMPG1 peptides, while green arrows indicate non-proteolyzed IMPG1. All western blots were replicated at least three times in different samples; total protein staining was used as a loading control for all repeated experiments (Fig. S1).

To exclude the possibility that sample deglycosylation leads to IMPG1 proteolysis, we examined IMPG1 Rf in serial dilutions of the protein deglycosylation mix (Fig. 2C). The results show that IMPG1 molecular weight increased in distinct stages proportional to enzyme dilutions and always appeared as a single band. In other words, serial dilution revealed linear increments of glycosylation on IMPG1 and not a binary state of the protein between proteolyzed and non-proteolyzed. These results support the premise that the deglycosylation treatment does not inadvertently cause IMPG1 proteolysis. In addition, ChSabc treatment was scrutinized as a possible cause of IMPG1 proteolysis. Molecular weight of IMPG1 was assessed in retinas of a mouse model constitutively lacking ChS, known as “small with kinky tail” (SKT) mice (Fig. 2D). SKT retinas showed an IMPG1 profile similar to WT mice, indicating that IMPG1 proteolysis was not due to ChSabc treatment. Moreover, these findings demonstrate that IMPG1 proteolysis is independent of ChS attachments to the molecule.

Our previous work demonstrated that localization of IMPG1 depends on the presence of IMPG2 ^7^. To assess the influence of IMPG2 on IMPG1 proteolysis, we analyzed the molecular weight of IMPG1 in the retinas of IMPG2-KO mice (Fig. 2E). Results revealed similar bands and distributions as WT, indicating that IMPG1 proteolysis is independent of IMPG2.

Finally, we expressed recombinant IMPG1 with a C-terminal DDK epitope tag in HEK293 cells. Immunoblots of cell samples stained with a DDK-specific antibody and IMPG1 antibody showed two proteolyzed IMPG1 bands at ∼75 kDa (anti-IMPG1) and ∼24 kDa (anti-DDK) (Fig. 2F), indicating proteolysis of the expressed IMPG1 transgene. Moreover, non-proteolyzed ∼100-kDa bands were also detected. Overall, these results indicate that IMPG1 undergoes proteolysis as a part of its normal maturation in the retina and in tissue culture cells. Moreover, this process occurs independently of the presence of IMPG2 and ChS.

### IMPG1 proteolysis and vision disease

Proteolytic SEA domains have a characteristic globular shape constituted by four alpha helices and four beta sheets, in which proteolysis occurs between the second and third beta sheets (Fig. 3A, B) ^14^. In humans, mutations affecting the SEA-2 domain of IMPG1 are linked to development of RP (Fig. 3A) ^11^. To determine if these mutations affect proteolysis of the SEA-2 domain, in HEK293 cells we expressed two *IMPG1* transgenes replicating mutations in the SEA-2 domain associated with RP. Western blot showed that cells expressing IMPG1-WT plasmid presented 72% proteolyzed and 28% non-proteolyzed protein, whereas cells expressing IMPG1-L583P or IMPG1-L630F presented 92% or 58% non-proteolyzed IMPG1, respectively (Fig. 3C, D). These results suggest that human RP vision loss associated with IMPG1 SEA-2 mutations may be partially due to deficient proteolysis.

**Figure 3.**
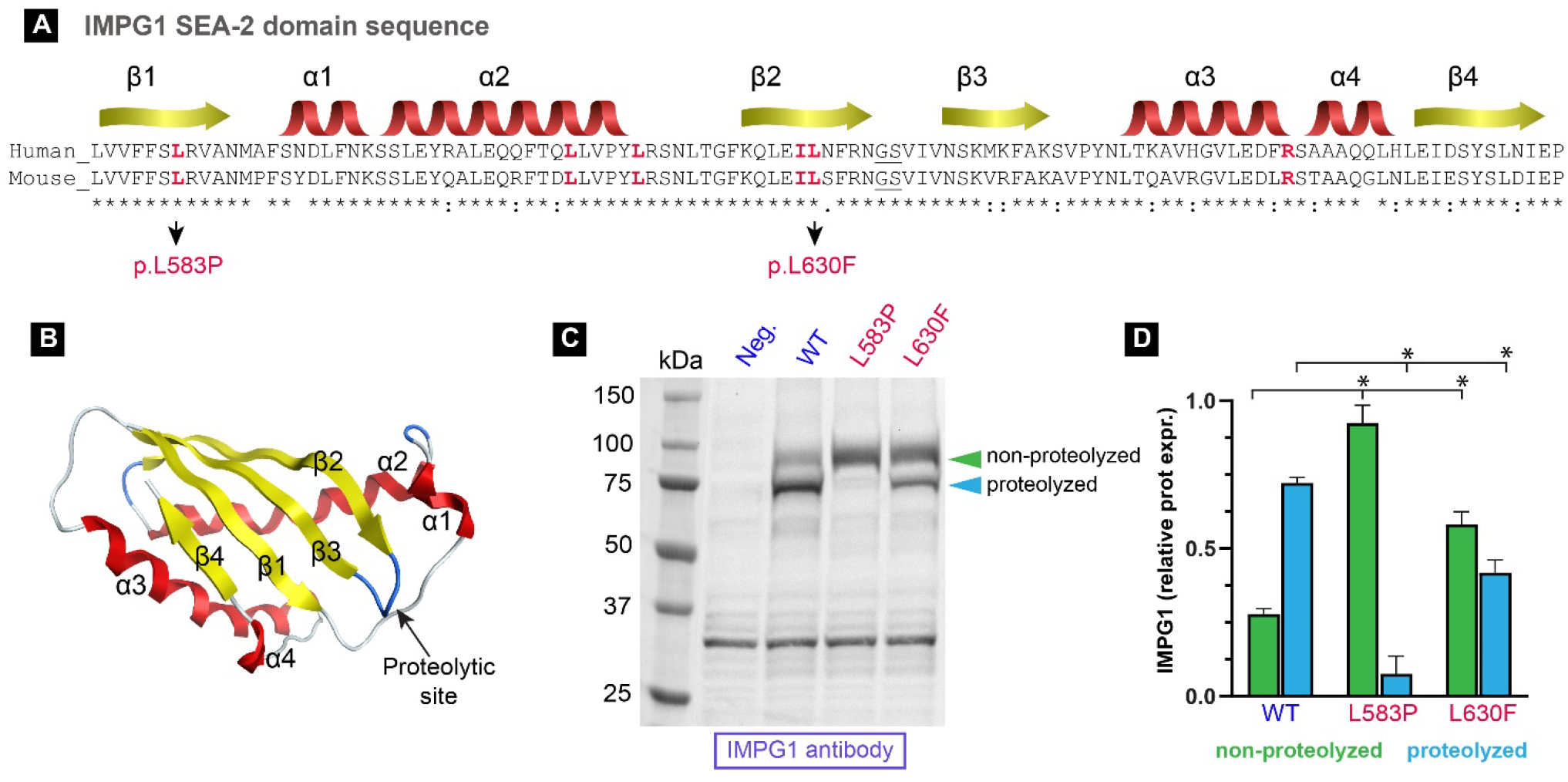
Mutations in the SEA-2 domain of IMPG1 prevent proteolysis. **A, B**, IMPG1 SEA-2 amino acid sequence and predicted tertiary structure, including mutations linked to retinitis pigmentosa pathology in humans (red letters). Protein structure prediction made with the AlphaFold server. **C**, Western blot assay detecting IMPG1 molecules synthesized by HEK293 cells expressing either GFP as a negative control, IMPG1-WT, IMPG1-L583P, or IMPG1-L630F transgenes. **D**, Protein expression quantification of proteolyzed (blue) and non-proteolyzed (green) IMPG1. Results were obtained in triplicate and referenced with total protein stain. Bars are mean ± SEM of three samples. Both WT groups are significantly different (*p<0.01) from each mutant by two-way T-test. All western blot samples were deglycosylated before loading onto the gel; untreated protein and total protein are in Figure S2.

### Proteolysis and translocation of IMPG2

The transmembrane proteoglycan IMPG2 has a theoretical molecular weight of 138 kDa, and SEA-2-mediated proteolysis is expected to liberate a 106-kDa portion of the extracellular region from a membrane-retained 32-kDa fragment (Fig. 4A). Western blot comparing WT and IMPG2-KO retinas using an antibody against the theoretical membrane portion of IMPG2 (IMPG2m) revealed a single band at 32 kDa with no additional bands at higher molecular weight in WT retinas (Fig. 4B). This result supports the hypothesized proteolysis of IMPG2. Further, retina lysates from mice lacking IMPG1 or ChS suggested that proteolysis of IMPG2 does not depend on the presence of IMPG1 or ChS. Antibody targeting the 106-KDa extracellular portion of IMPG2 could not detect IMPG2 by western blot, despite sample treatments with Deglyco and ChSabc enzymes [not shown; loading controls in supporting information (Fig. S3)].

**Figure 4.**
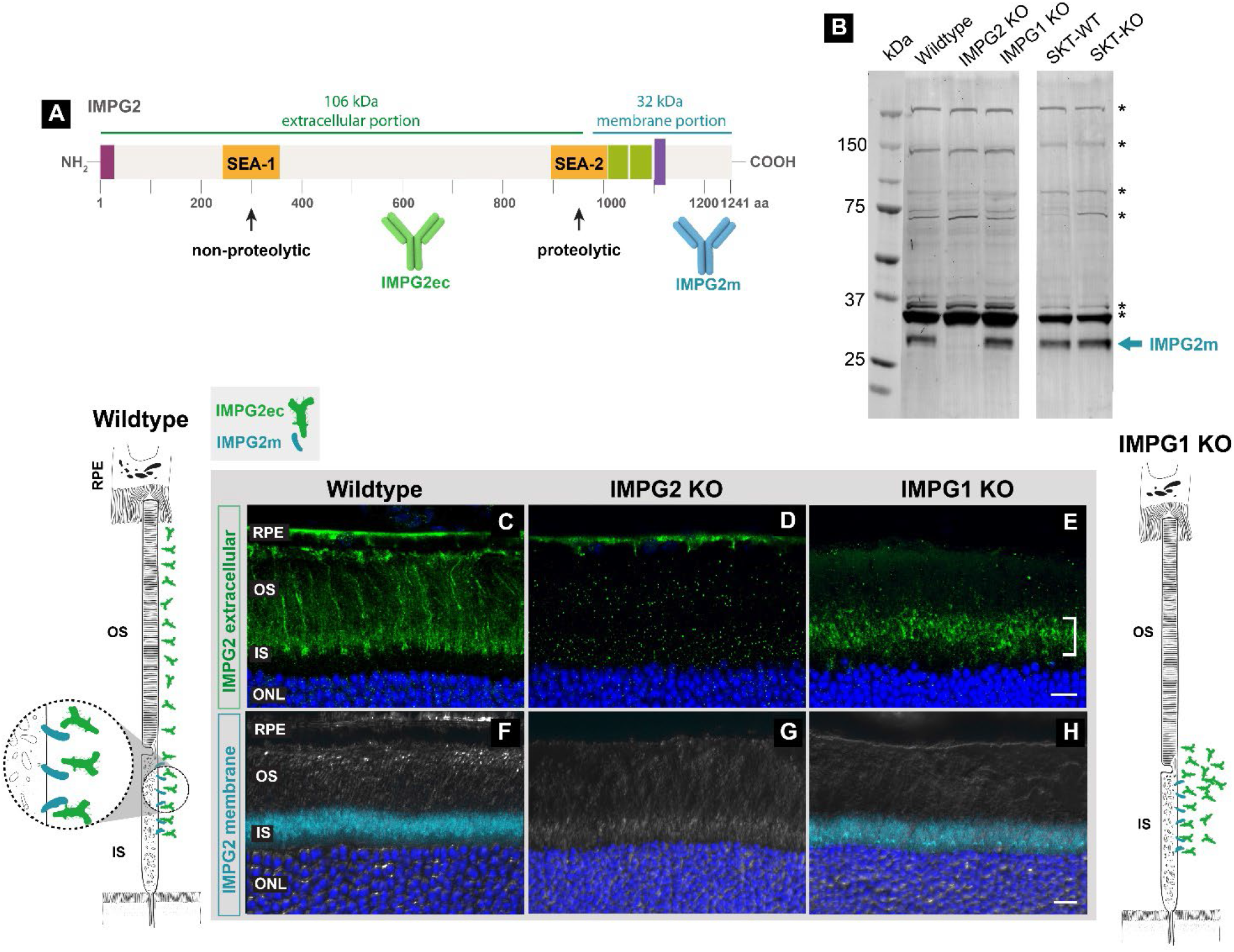
IMPG2 proteolysis and traffic of its extracellular portion from the matrix inner segment (IS) to outer segment (OS) in an IMPG1-dependent process. **A**, Schematic of IMPG2 shows the expected proteolytic and non-proteolytic SEA domains along with extracellular and membrane-portion antibody recognition sites. **B**, Representative western blot comparing retina samples from wildtype (WT), IMPG2 knockout (KO), IMPG1 KO, and SKT mice WT and KO. A single band at 32 kDa is present in wildtype, IMPG1 KO, and SKT mice (cyan arrow). **C–H**, Immunohistochemistry staining on retina cross-section from wildtype (**C, F**), IMPG2 KO (**D, G**), and IMPG1 KO (**E, H**) mice using a specific antibody against the extracellular portion of IMPG2 (IMPG2ec, green) and membrane portion (IMPG2m, cyan). IMPG2ec antibody stains the outer part of the IS and entire OS (**C**). IMPG2ec localizes at the outer IS and inner OS region (white bracket), with a substantial reduction in the rest of the OS (**E**). Outer nuclear layer (ONL) stained with DAPI (blue). Differential interference contrast (DIC) in white. RPE indicates retina pigment epithelium. Scale bar equals 10 µm. N=3 mice per experiment at postnatal day 45. Scheme of IMPG2m and IMPG2ec localization in wildtype and IMPG1 KO mice according to immunohistochemistry results (**C, F, E, H**).

We previously described localization of IMPG2 at the photoreceptor IS using a primary antibody recognizing the membrane portion of IMPG2 (IMPG2m) ^7^. Considering IMPG2 proteolysis, the extracellular portion of IMPG2 (IMPG2ec) may not co-localize with IMPG2m peptide. Immunohistochemistry analysis of WT retinas using antibodies targeting IMPG2ec revealed that it localizes around the IS and OS of photoreceptors, while IMPG2m is localized to the IS (Fig. 4C, F). These key findings demonstrate that IMPG2 undergoes intramolecular proteolysis and generates an extracellular portion that travels from the inner IPM to the outer IPM.

Past work indicates that retinas from mice lacking IMPG2 have aggregation and mislocalization of IMPG1. However, lack of IMPG1 does not affect localization of IMPG2m (Fig. 4H) and unknown effects on IMPG2ec ^7^. To determine the role of IMPG1 in IMPG2ec localization, we examined localization of IMPG2ec in IMPG1-KO retinas. Immunohistochemistry analysis revealed that IMPG2ec mislocalized at the intersection between IS and OS, leaving the rest of the OS depleted of IMPG2ec in mice lacking IMPG1 (Fig. 4E). These results indicate that localization of IMPG2ec depends on the presence of IMPG1, similar to the localization dependence of IMPG1 on IMPG2 expression ^7^. Together, these results imply reciprocal dependency for proper photoreceptor localization between IMPG1 and IMPG2.

## Discussion

Our results indicate that both IMPG1 and IMPG2 undergo intramolecular proteolysis in the SEA-2 domain as part of the protein maturation process. This work also shows that mutations in the SEA-2 domain of IMPG1 associated with RP human vision disease affect IMPG1 proteolysis. Importantly, our work along with previous results ^7^ indicate that IMPG1 and IMPG2 traffic from the IS matrix to the OS matrix through a mechanism that requires presence of both IMPG1 and IMPG2.

SEA domains are associated with secreted and glycosylated membrane proteins and can be either proteolytic or non-proteolytic ^9,13^. IMPG1 and IMPG2 possess two SEA domains in their molecule structure: SEA-1, which is a non-proteolytic domain and SEA-2, a proteolytic domain (Fig. 1A) ^6,10,19^.

Our analysis predicts that SEA-1 domains in IMPG1 and IMPG2 have amino acid sequences that do not match known proteolytic sequences (Fig. 1B). Our data support this prediction, given that hypothetical proteolysis in IMPG1 at SEA-1 would generate a peptide with a lower molecular weight than observed in our assays. For IMPG2, we were not able to test the non-proteolytic prediction in SEA-1 since we could not analyze the extracellular IMPG2 peptide by western blot because IMPG2ec antibody does not work in western blot. The function of SEA-1 domains in IMPG molecules remains to be tested in future studies.

Our results indicate that IMPG1 proteolyzes independently of the presence of IMPG2 or ChS. In humans, IMPG1 point mutations within the SEA-2 domain generate photoreceptor degeneration linked to RP, whereas mutations outside SEA-2 lead to vitelliform macular dystrophy ^11,20,21^. This demarcation between mutations linked to RP or vitelliform macular dystrophy in IMPG1 is not mirrored in IMPG2, where mutations within and outside the SEA domains can generate either pathology ^22-26^. Our results showed that two mutations in IMPG1 linked to RP obstruct IMPG1 proteolysis in vitro. These results suggest that deficits in IMPG1 proteolysis are involved in the pathophysiology of RP linked to mutations in IMPG1.

The potential for IMPG2 to undergo SEA-mediated proteolysis has been previously overlooked. Here we showed that IMPG2 undergoes cleavage at the SEA-2 domain to generate extracellular and membrane-bound peptides. Our localization studies indicated that IMPG2ec is released from the IS to distribute evenly between the IS and OS. These results help refine our understanding of the parameters governing IMPG localization (Fig. 5). In this model, both IMPG molecules are synthesized at the photoreceptor IS and secreted to the IPM adjacent to the IS. IMPG1 and IMPG2ec then traffic to the IPM around the OS using an undetermined mechanism. Further, our results show that trafficking of IMPG2ec depends on the presence of IMPG1. Likewise, we previously demonstrated that proper localization of IMPG1 relies on the presence of IMPG2 ^7^. Together these results indicate that both IMPG molecules are co-dependent for their final localization around the OS.

**Figure 5.**
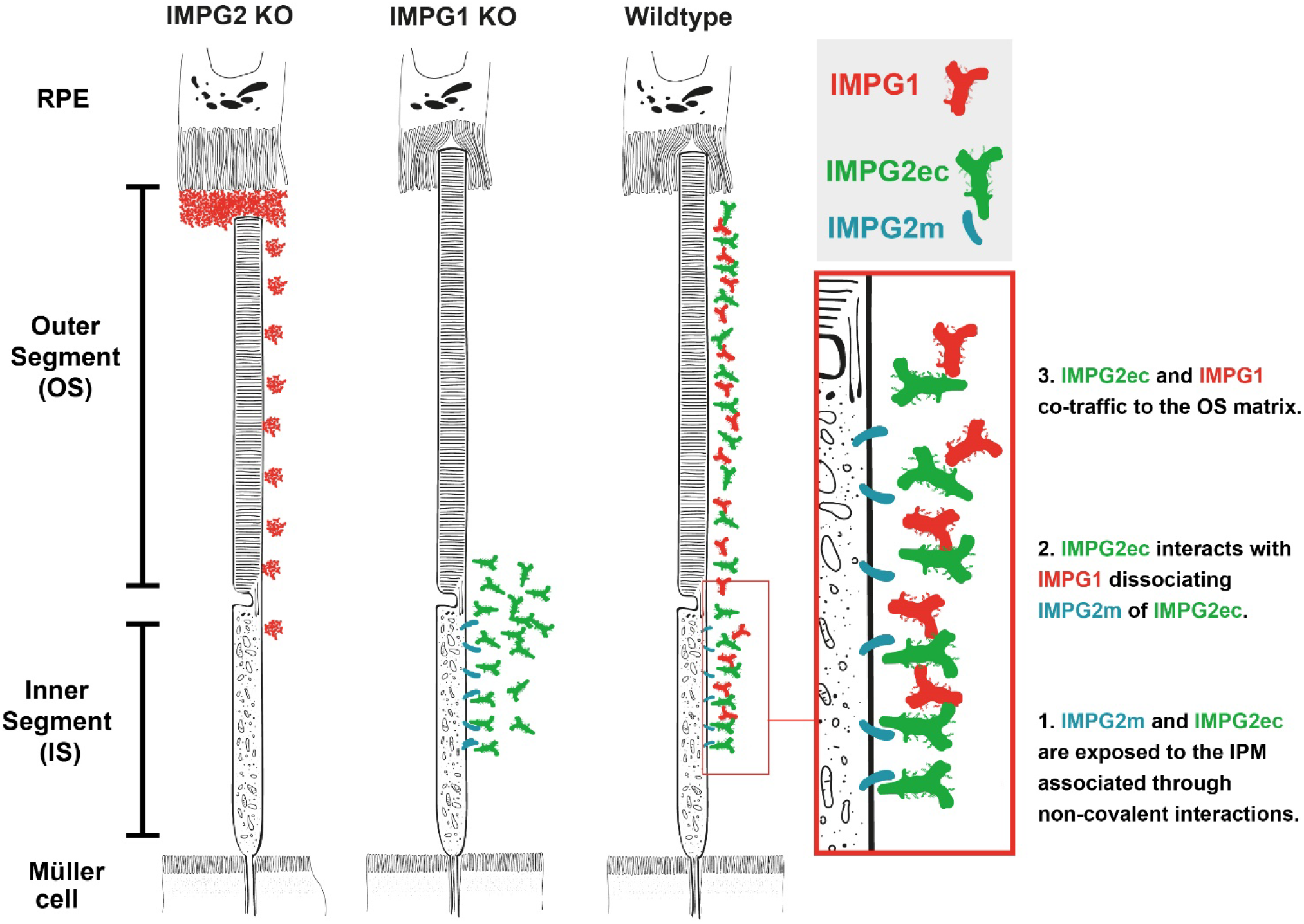
Schematic of IMPG1 and IMPG2 retina localization in mice lacking IMPG2, IMPG1, and wildtype after integration of findings from our study. **Left panel** represents IMPG2 knockout (KO) mice retina showing IMPG1 (red) aggregation and accumulation at the intersection between retina pigment epithelium (RPE) and the photoreceptor outer segment (OS), as described in our previous work. **Center panel** illustrates IMPG1 KO retina showing mislocalization of extracellular IMPG2 (IMPG2ec, green) around the inner OS and outer inner segment (IS). IMPG2 membrane peptide (IMPG2m, cyan) maintains it IS localization. **Right panel** illustrates current understanding of IMPG2 localization, where the IMPG2m anchors to the IS cell membrane, and IMPG2ec localizes at the IS and OS interphotoreceptor matrix (IPM). **Red box** shows a magnified view of the IS with a speculative representation of IMPG2m and IMPG2ec dissociation after interaction with proteolyzed IMPG1 at the IS to further traffic towards the OS.

Cell culture assays expressing mucin-3 proteoglycan containing a SEA domain demonstrate that proteolytic cleavage occurs in the endoplasmic reticulum ^27^. After proteolysis, the generated subunits recognize each other to form a heterodimeric complex through a non-covalent SDS-sensitive interaction ^14,16,27,28^. Proteolysis of IMPG molecules generates SEA “sticky” ends with the potential to associate through non-covalent interactions ^9,28^. This speculative scenario lets us hypothesize about generation of heterodimers between IMPG1 and IMPG2 subunits as part of a hypothetical IMPG1–IMPG2 complex assembly needed to traffic and distribution of IMPG proteins into the OS region. Specifically, we hypothesize that non-covalent interactions between IMPG2m and IMPG2ec subunits occur at the photoreceptor IS, and they rapidly detach after being exposed to the extracellular space containing proteolyzed IMPG1, forming an IMPG1–IMPG2 complex needed to traffic towards the IPM around the OS (Fig. 5, red rectangle).

In summary, these results show that proteolysis is part of the maturation process for both IMPG molecules and suggest that mutations in IMPG1 linked to vision disease in humans affect proteolysis. Further, our results show mutual localization dependency between IMPG1 and IMPG2 to traffic from the IPM around the IS to the OS. These results indicate the existence of an undescribed trafficking system and interaction between IMPG1 and IMPG2, where the affection of one protein affects the localization of the other protein. This work proposes that mutations in IMPG1 or IMPG2 leading to vision diseases could be caused by defective localization of both mutant and normal-partner IMPG proteins.

## Materials and methods

### Animals

IMPG1 KO and IMPG2 KO animal models were generated at the West Virginia University transgenic core with the use of Crispr-Cas9, as described in Salido et al. (2020) ^7^. SKT (small with kinky tail) mice, also known as CHSY1 KO mice, were purchased from Jackson Laboratory (stock no: 001433). All mouse populations were backcrossed with C57BL/6J (Jackson Laboratory) and consistently genotyped to identify possible *RD1* and *RD10* mutations. Animals were maintained under 12-h light/dark cycles with food and water ad libitum. Male and female mice were equally used throughout experiments. All experimental procedures involving animals were approved by the Institutional Animal Care and Use Committee of West Virginia University and all experiments were performed in accordance with relevant guidelines and regulations. The manuscript follows the recommendations in the ARRIVE guidelines.

### Western blot

Mice were euthanized using CO2, followed by cervical dislocation as a secondary method. Eyes were enucleated, corneas and lenses surgically removed, and retinas carefully extracted from the eyecup and rapidly frozen on dry ice. Frozen retinal samples were sonicated in T-PER buffer (Thermo Fisher, cat. 78510) with HALT protease and phosphatase inhibitor/EDTA (Thermo Fisher, cat. 78430) as recommended. Protein concentrations were measured using a standard BCA protein assay (Pierce, cat. 23225). Samples treated with chondroitinase ABC (ChSabc) (Sigma, cat. C3667) were incubated for 3 hours at 37°C with 0.1 units per sample, with periodic vortex homogenization every hour. Likewise, deglycosylation treatment (Dglyco) consisted of sample incubation for 3 hours at 37°C with 20 nL of Protein Deglycosylation Mix II (NEB, cat. P6044S) per μL of sample. Immediately following enzyme treatment, LDS buffer (Genscript, cat. M00676) mixed with B-mercaptoethanol was added to the samples and stored at -20°C.

Equal sample concentrations (100 μg of total protein per well) were resolved in 4%–20% electrophoresis gradient gels (Genscript, Cat. M00656) and transferred onto polyvinylidene difluoride (PVDF) membranes (Immobilon-FL; Millipore). Membranes were stained with Total Protein Staining kit (LI-CORE, cat. 926-11010), imaged at 700 nm, and immediately blocked (Thermo Fisher, cat. 37570) for 1 hour at room temperature. Membranes were incubated overnight at 4°C on a bidirectional rocker with IMPG1 and IMPG2m primary antibodies, as previously described ^7^. Following primary antibody incubation, membranes were washed in PBST (PBS with 0.1% Tween-20) three times for 5 minutes each at room temperature. Secondary antibody incubation was conducted for 45 minutes at room temperature using goat anti-mouse Alexa Fluor 680 (Invitrogen, cat. A28183) or donkey anti-mouse IRDye 800 (Li-Cor, cat. 926-32212) and goat anti-rabbit Alexa Fluor 488 (Invitrogen, cat. A11034). Membranes were washed three times for 5 minutes each with PBST and scanned using a GE Typhoon 9410 imager.

### Plasmid design and protein expression

Human IMPG1 with a DDK tag in the C-terminus, under the CMV promotor in an MR210410 vector, was generated by Origene. The second set of plasmids contained three versions of mouse IMPG1: one WT, and two with point mutations in L583P or L630F. Mutations were confirmed by sequencing. Full-length mouse IMPG1 variants were synthesized and inserted in a pcDNA3.1(+)-P2A-eGFP vector under a CMV promotor by Genscript. All IMPG1 proteins were expressed in HEK293 cells. Cells were collected 48 hours post-transfection and sonicated in PBS with protease inhibitor mixture (Roche). Samples were processed and analyzed by western blot as described above.

### Immunohistochemistry

All mice were euthanized with CO2 and cervical dislocation, after which eyes were carefully enucleated. A small hole was cut into each cornea, and enucleated eyes were placed in 3% paraformaldehyde for 3 hours at room temperature. Increasing serial dilutions of 7.5%, 15%, and 20% sucrose solution were interchanged over a two-day period. Eyecups were placed in optimal cutting temperature compound (Sakura) and flash-frozen in an alcohol bath on dry ice prior to storage at -80°C.

Using a Leica CM1850 cryostat, 16-μm cross-sections were cut and carefully placed on Superfrost Plus slides (Fisher Scientific). Immunofluorescent staining and image acquisition were concurrently performed as described in past studies ^7^, with addition of the following treatment. To specifically target the extracellular region of IMPG2, two enzyme digestion steps were required before proceeding with standard immunostaining: 1) 1 U/mL of ChSabc enzyme (Sigma, cat. C3667) was added to the slide and incubated for 1 hour at 37°C; 2) 7 µL of Protein Deglycosylation Mix II (NEB, cat. P6044S) per slide was added for deglycosylation and incubated overnight at 37°C. After both assays, the standard immunostaining technique was used as previously described ^7^. We used a primary antibody against the extracellular region of IMPG2 (Sigma, HPA015907) at 1:500 dilution.

Images were acquired and processed with a Nikon C2 laser-scanning confocal microscope using differential interference contrast (DIC) and excitation wavelengths of 405, 488, 543, and 647 nm with 60X objectives. Sections scanned in Z where 10 µm deep in 0.2 µm steps and merged by maximum intensity projection. The lookup table (LUT) is linear and covers the full range of the data. Four sections were imaged for each sample at room temperature, and data were derived from three independent experiments using littermates as controls. Adobe Photoshop and Illustrator software were used to crop images and create the final figures.

### Experimental design

All mice used in this work were postnatal day 45 at the time of euthanasia. Two retinas from the same mouse were collected in one tube for western blot experiments. All results were reproduced at least three times using samples from different mice. For immunohistochemical analysis, at least four sections were imaged per sample, and data were derived from three independent experiments. Sex differences were assessed for each outcome measured, with no significant variance observed.

## Data availability statement

This article contains supporting information. No datasets were generated or analyzed during the current study.

## Author contributions statement

BM: Investigation, Resources, Writing- Reviewing and Editing; CC: Investigation, Writing- Reviewing and Editing; WJG: Resources, Writing- Reviewing and Editing; SR: Resources; EMS: Conceptualization, Methodology, Writing - Original Draft, Supervision, Project administration, Funding acquisition.

## Funding and Acknowledgements

This work was supported by Knights Templar Eye Foundation, West Virginia Lions and Lions Club International Foundation, Visual Sciences CoBRE (grant P20GM144230), and West Virginia University. We thank Visvanathan Ramamurthy, PhD, Mario Ermacora, PhD, and Scott Weed, PhD, for discussion of results, as well as Lucia Bonifacini for helping prepare figures and WVU Research and Graduate Education for manuscript editing. We also thank Peter Mathers, PhD, and the WVU transgenic core for their roles in the generation of our animal models, which is funded in part by the WV Tumor Microenvironment CoBRE (grant P20GM121322).

