## Supplementary Information for "Interphotoreceptor matrix proteoglycans IMPG1 and IMPG2 proteolyze in the SEA domain and reveal localization mutual dependency"

Supplementary Figure S1
Supplementary Figure S2
Supplementary Figure S3

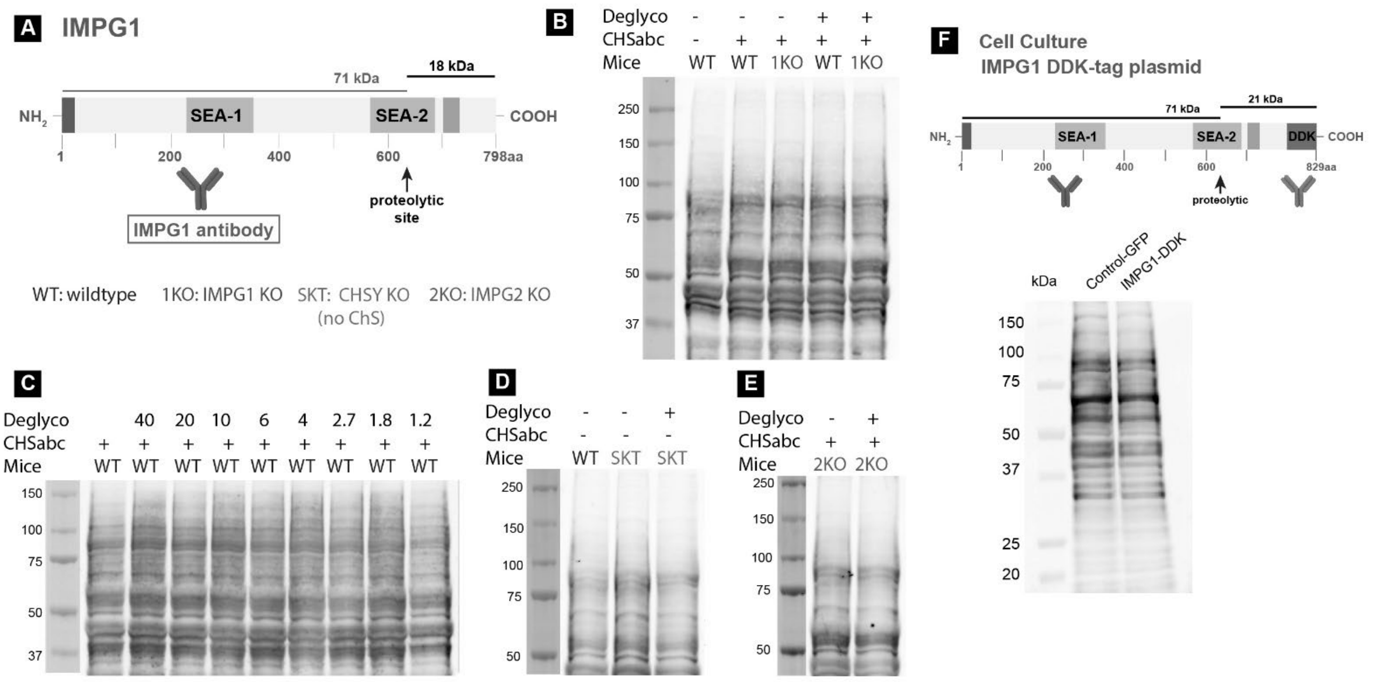


**Supplementary Figure S1**. Western blot loading control of Figure 2 by total protein staining assay.

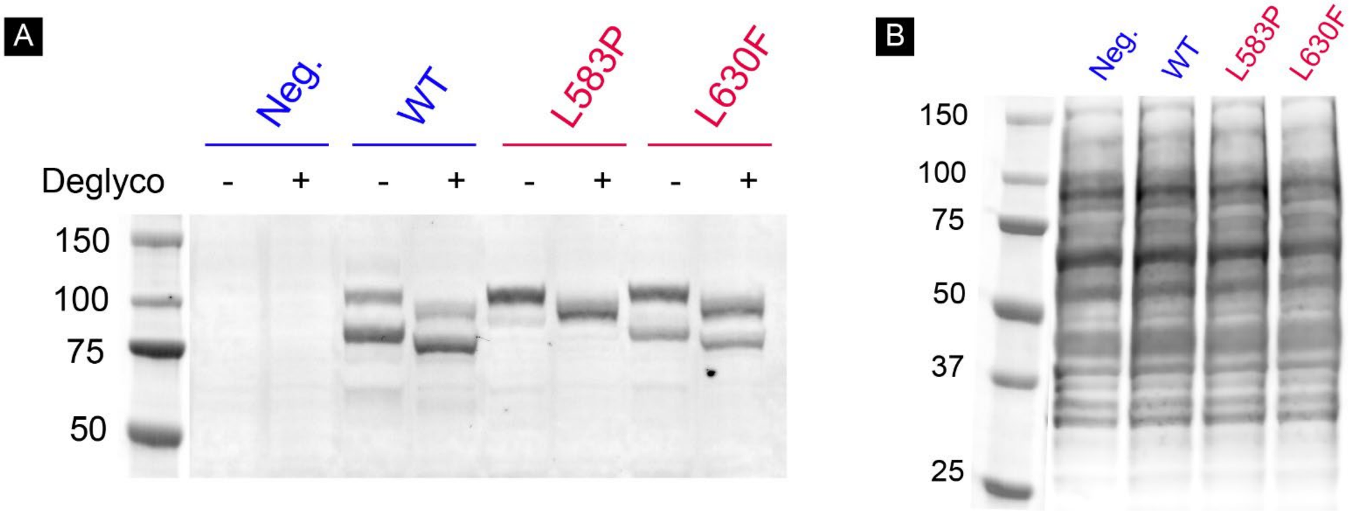


**Supplementary Figure S2**. **HEK293 cells glycosylate IMPG1 unrelatedly to protein proteolysis. A**, the same samples used in figure 3, IMPG1 WT and mutated proteins expressed in HEK293 cells treated with or without deglycosylation protein mix (Deglyco). Glycosylation in IMPG1 generates a ~5kDa increase in the relative molecular mobility in proteolyzed and non-proteolyzed molecules. **B**, Western blot loading control of Figure 3 by total protein staining assay. The experiment was reproduced 3 times with different samples.

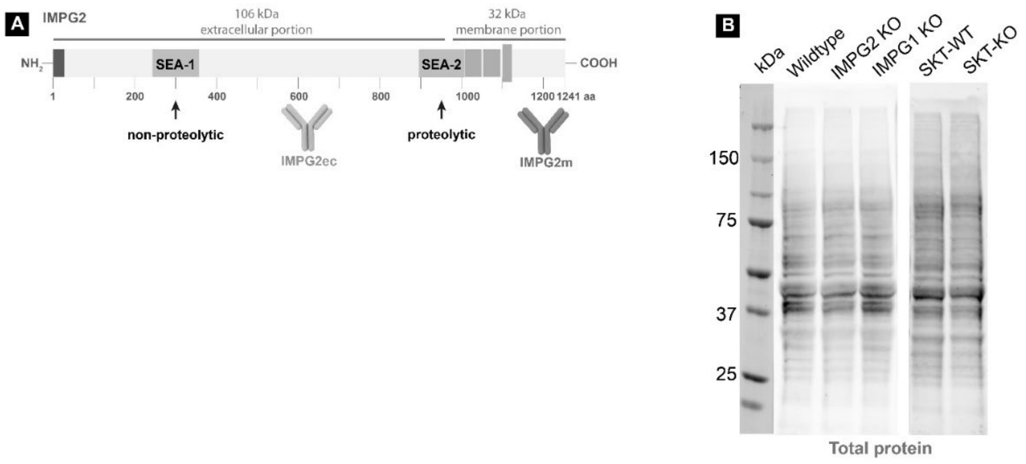


**Supplementary Figure S3**. Western blot loading control of Figure 3b in the text of the manuscript by total protein staining assay.
